# Antioxidant capacity of Catechinopyranocyanidins derived from adzuki beans

**DOI:** 10.64898/2026.05.15.725587

**Authors:** Riki Kawabata, Ikki Hagiwara, Nene Komizo, Yuto Inaba, Takanori Matsui, Takashi Ito

## Abstract

Catechinopyranocyanidins (Cpcs) which consist of diastereomers A and B are pigments derived from adzuki beans and are compounds in which the catechin and cyanidin skeletons are condensed to a pyrano ring. While catechins and anthocyanidins possess high antioxidant capacity, the physiological functions of Cpcs remains unclear. In this study, the antioxidant capacity of Cpcs was evaluated by in vitro antioxidant assays and by assessing their cytoprotective activity against oxidative stress in normal human dermal fibroblasts (NHDFs). Antioxidant capacity based on the hydrogen atom transfer (HAT) mechanism, as assessed by the ORAC assay revealed that Cpcs exhibit 14.1 μmol TE/μmol (Trolox equivalent antioxidant capacity: TEAC). Meanwhile, capacity based on the single electron transfer (SET) mechanism, as assessed by the DPPH, ABTS and CUPRAC assays revealed, they exhibit 2.1–3.6 μmol TE/μmol. Since TEAC value of Cpcs demonstrated by the HAT based mechanism higher than its SET based oxidative capacity suggesting that the antioxidant capacity of Cpcs is driven by the HAT mechanism. In cell culture experiments, Cpcs ameliorate cell toxicity in rotenone-induced injury model, suggesting to cytoprotective activity against mitochondrial dysfunction-dependent apoptosis. These results reveal novel physiological functions of Cpcs which may serve as a design guideline for elucidating in vivo dynamics based on antioxidant mechanisms.

## Introduction

The oxidative stress response in human body plays a crucial role in regulating chronic inflammation, cellular senescence and cell death, and a decline in its activity trigger age-related diseases [1-3]. Therefore, intervention with the antioxidant factors derived from foods is expected to contribute to health promotion. For instance, Flavan-3-ols, such as catechin possess strong radical scavenging capacity [4–6] and have been shown to potentially alleviate conditions such as cardiovascular disease, obesity, and diabetes [7–9]. Similarly, anthocyanidins, such as cyanidin, also exhibit antioxidant activity [10–12], and they may be effective in alleviating diseases such as intestinal inflammation and atherosclerosis [13–14].

Adzuki bean (*Vigna angularis*) is legume native to Southeast Asia and first domesticated in Japan as “adzuki bean” [15]. In Japan, small red beans have long been rooted in culture, with various uses explored. While numerous cooking methods have been developed for their consumption, adzuki has attracted considerable attention due to their rich nutritional content recently [16–19]. Defining characteristics of adzuki is their vivid reddish-purple color. Recently, Yoshida et al have identified catechinopyranocyanidins A and B (Cpcs) as pigment of adzuki [20–22], which are hydrophobic compounds formed by the combination of a cyanidin skeleton with a catechin skeleton, creating a pyrano ring. The A and B types are distinguished by differences in the stereochemical configuration of the catechin skeleton portion [21]. Given that Cpcs contain catechin and cyanidin skeletons, it is plausible that Cpcs may exhibit strong antioxidant properties contribute to adzuki’s health benefits. However, the antioxidant capacity of Cpcs have not yet been reported.

In the present study, we evaluated the antioxidant capacity of Cpcs *in vitro* and further examined its cytoprotective effects against oxidative stress in mammalian cells.

## Materials and methods

### Isolation of catechinopyranocyanidins from adzuki beans

Cpcs were isolated from adzuki beans as previously reported with slight modification [21]. Briefly, adzuki beans (*Vigna angularis* cv. Erimoshozu; Mamehei, Toyama, Japan) were boiled and seed coat was isolated. Isolated seed coat was soaked in of ethyl acetate for 24 h, and this process was repeated three times. The extract was collected, evaporated to dryness, and redissolved in 5 mL of methanol to obtain the crude extract. Purification of Cpcs was performed in two steps. First, gel filtration chromatography was conducted using a chromatography column (20 I.D., 300 mm; Climbing, Fukuoka, Japan) and TOYOPEARL HW-40C resin gel (Tosoh, Japan), pre-equilibrated with 100 mL of methanol. The extract was applied and eluted stepwise with 10% acetone in methanol, followed by increasing acetone concentrations in 10% increments. The purple fractions were collected and evaporated to dryness to obtain crude Cpcs. Second, the crude Cpcs were dissolved in 2 mL of methanol, filtered through a 0.45 μm membrane, and subjected to high-performance liquid chromatography (DIONEX UltiMate 3000 HPLC system; Thermo Fisher Scientific, Waltham, MA, USA) equipped with a FLEX FIRE C30 column (10.0 I.D., 250 mm; Nomura Chemical, Aichi, Japan). The mobile phase consisted of 0.5% trifluoroacetic acid in water (A) and acetonitrile containing 0.5% trifluoroacetic acid (B). Elution was monitored at 280 and 570 nm using a photodiode array detector. The column temperature was maintained at 40°C, and the injection volume was 80 μL. The collected fractions of Cpcs (15 fractions) were evaporated to dryness, and the concentration was calculated from the dry weight obtained, and the sample was dissolved in methanol to determine the absorbance and molar absorption coefficient.

### Antioxidant capacity evaluation

For all antioxidant capacity evaluation tests, the assays were performed according to previous studies [23–25] with some modifications. In DPPH and ABTS assay, calibration curves were constructed using Trolox (Tokyo Chemical Industry, Japan) as the standard compound, based on the relationship between Trolox concentration and radical scavenging rate calculated as follows: Scavenging rate (%) = (*Abs*_*sample*_ ― *Abs*_*blank*_ /*Abs*_*control*_ ― *Abs*_*blank*_) × 100. Samples were prepared at appropriate concentrations, and only values falling within the linear range of the Trolox standard curve were used for analysis. The Trolox equivalent concentration (μM TE) obtained from the calibration curve was divided by the original sample concentration (μM) to calculate the Trolox equivalent antioxidant capacity (TEAC, μmol TE/μmol).

### Diphenyl-1-picrylhydrazyl (DPPH) assay

190 μL of 0.1 mM 2,2-diphenyl-1-picrylhydrazyl (DPPH; Nacalai Tesque) methanol solution was mixed with 10 μL of sample in a 96-well plate and incubated at 37°C for 30 min. Methanol was used as a control, and a blank without DPPH was included. Absorbance was measured at 518 nm. The EC_50_ (μM) was defined as the concentration required to achieve 50% radical scavenging rate.

### 2,2′-azinobis(3-ethylbenzothiazoline-6-sulfonic acid) (ABTS) assay

ABTS radical cations were generated by reacting 7 mM 2,2′-azinobis(3-ethylbenzothiazoline-6-sulfonic acid)diammonium salt (ABTS; Nacalai Tesque) with 140 mM potassium peroxydisulfate (Nacalai Tesque) in the dark at room temperature for 16 h, and diluted with methanol to an absorbance of 0.700 ± 0.020 at 734 nm. 190 μL of resulting solution was mixed with 10 μL of sample in a 96-well plate and incubated at 37°C for 4 min. Methanol was used as a control, and a blank without ABTS was included. Absorbance was measured at 734 nm. The EC_50_ was calculated as described for the DPPH assay.

### Copper reduction antioxidant capacity (CUPRAC) assay

50 μL of 0.5 M ammonium acetate buffer (pH 7.2), 50 μL of 10 mM copper(II) sulfate pentahydrate solution, and 50 μL of 7.5 mM neocuproine (Tokyo Chemical Industry) solution in methanol were added to a 96-well plate, followed by 30 μL of ultrapure water and 20 μL of sample. The mixture was incubated in the dark at room temperature for 30 min. Ultrapure water was used for blank instead of the copper(II) sulfate solution. Calibration curve was constructed based on the relationship between absorbance at 450 nm and the corresponding Trolox concentration, and the reducing power of copper(II) ions was quantified and expressed as TEAC.

### Oxygen radical absorbance capacity (ORAC) assay

30 μL of 75 mM phosphate buffer (pH 7.4), 20 μL of sample, and 120 μL of 0.05 μM fluorescein (Nacalai Tesque) solution prepared in the same buffer were added to a 96-well black plate and incubated at 37°C for 30 min. Methanol was used for blank instead of the sample. Subsequently, 50 μL of 150 mM 2,2′-azobis(2-methylpropionamidine)dihydrochloride (AAPH; FUJIFILM Wako Pure Chemical, Osaka, Japan) prepared in 75 mM phosphate buffer was added. Fluorescence was recorded immediately (Ex 495 nm, Em 528 nm) every 5 min for 120 min at 37°C. The area under the fluorescence decay curve (AUC) was calculated, and the net AUC was obtained by subtracting the blank. ORAC values were calculated based on a calibration curve constructed from the relationship between net AUC and the corresponding Trolox concentration, and expressed as TEAC.

### Cell culture

Normal human dermal fibroblasts (NHDFs; Cat no: C-12302, lot: #435Z020.3, PromCell, Heidelberg, Germany) were cultured in Dulbecco’s modified Eagle’s medium (Nacarai Tesque) containing 10% fetal bovine serum (Biowest, Riverside, MO, USA) and 1% penicillin (100 U/mL) + streptomycin (100 μg/mL) + amphotericin B (0.25 μg/mL) (Nacarai Tesque) as previously described [26]. The culture conditions were set at 37°C and 5% CO_2_.

### Cell viability assay

NHDFs were cultured in 96-well plates and cultured with 1 or 10 μM of Cpcs for 24 h. After that, cell viability was measured using cell counting kit-8 (Dojindo, Kumamoto, Japan) according to the manufacturer’s protocols.

### Cytoprotective activity assay

NHDFs were cultured in 96-well plates and cultured with 10 μM of Cpcs for 24 h. After cultivate with Cpcs, cells were cultured with 1 μM of rotenone (Rot) for 24 h. Cytoprotective activity was measured as a cytotoxicity alteration using LDH assay kit-WST (Dojindo) according to the manufacturer’s protocols.

To evaluate cytosolic DNA fragmentation, cells cultured in the same protocol were stained with DAPI (Thermo Fisher Scientific), and the percentage of morphological abnormality (nuclear fragmentation) was calculated from 5 microscopic field per well, for total of 15 microscopic field of view per group (BZ-X810; Keyence, Japan).

### Statistical analysis

Statistical analysis was performed using GraphPad Prism 11 (ver. 11.0.0, GraphPad Software, Boston, MA, USA). Analysis was conducted by one-way ANOVA followed by Tukey’s multiple comparisons. The significance criterion was set at *p* < 0.05.

## Results

### Cpc exhibits high levels of antioxidant capacity in vivo

Antioxidant–radical reaction mechanisms are broadly classified into two categories: single electron transfer (SET) in which an electron is donated to reduce the radical and hydrogen atom transfer (HAT) in which a hydrogen atom is donated. The DPPH, ABTS, and CUPRAC assays are based on SET mechanisms, whereas the ORAC assay reflects a HAT-based mechanism [27]. In this study, dibutylhydroxytoluene (BHT), a well-established reductant, was used as the control.

In the DPPH assay, the EC_50_ for Cpcs was 396 μM and for BHT was 1131 μM, indicating that the EC50 of Cpcs was approximately 35% of that of BHT (Fig 1A). In the TEAC value of Cpcs exhibited 2.2770 Trolox equivalents per 1 μmol which is approximately 4.3-fold higher than that of BHT (Fig 1B).

**Fig 1.**
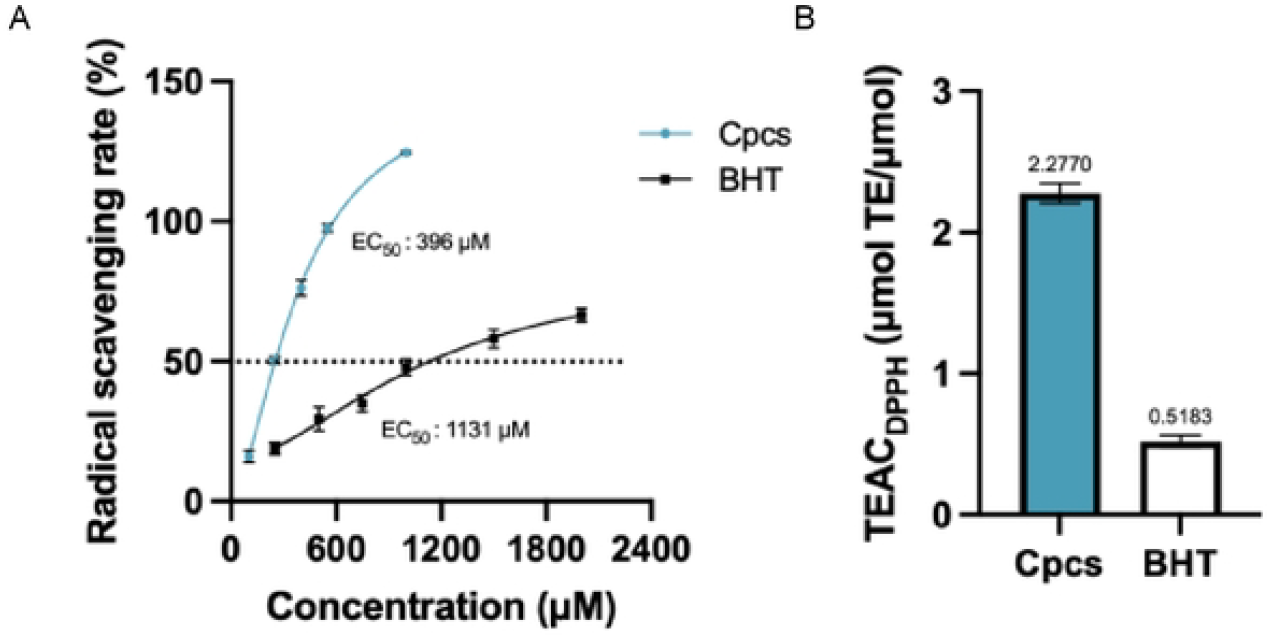
Comparisons of antioxidant capacity between Cpcs and BHT using the DPPH assay. (A) Radical scavenging rates calculated from the results of DPPH assay in 100–2000 μM of Cpcs and BHT are shown as sigmoidal curve. (B) The TEAC values of Cpcs and BHT calculated from the results of DPPH assay. Data are shown as mean ± SD. Similar results were obtained from other experiments.

In the ABTS assay, the EC_50_ for Cpcs was 253 μM and for BHT was 736 μM, indicating that the EC50 of Cpcs was approximately 34% of that of BHT (Fig 2A). In the TEAC value of Cpcs exhibited 2.1450 Trolox equivalents per 1 μmol, approximately 2.8-fold higher than that of BHT (Fig 2B).

**Fig 2.**
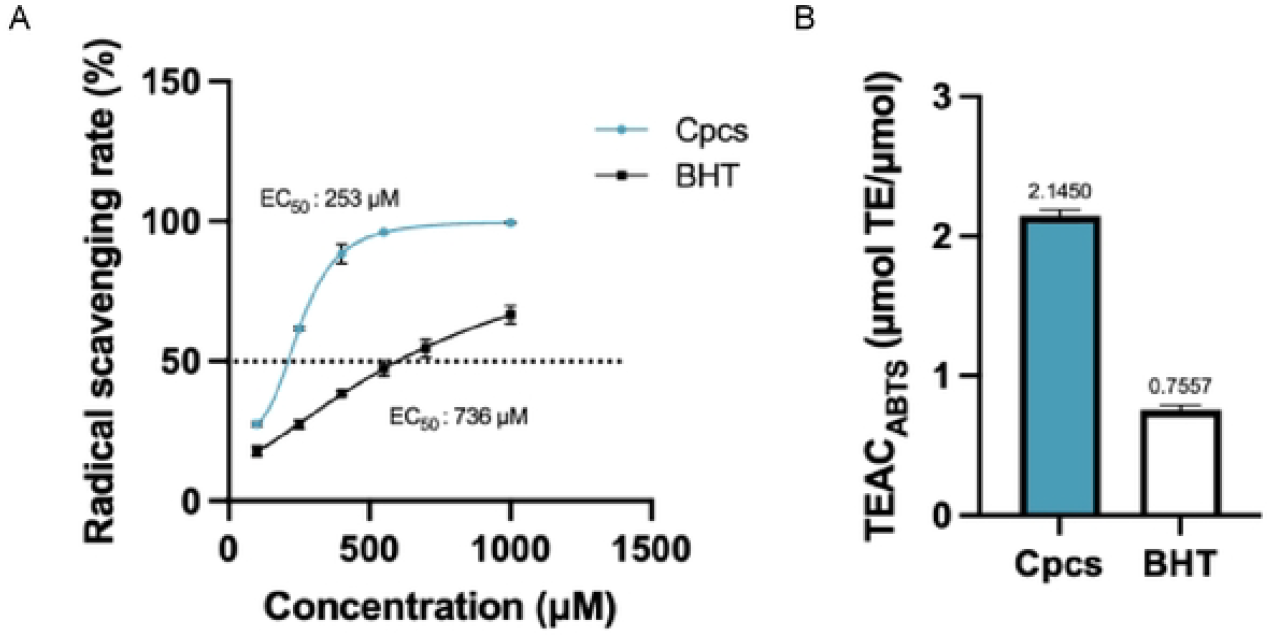
Comparisons of antioxidant capacity between Cpcs and BHT using the ABTS assay. (A) Radical scavenging rates calculated from the results of ABTS assay in 100–1000 μM of Cpcs and BHT are shown as sigmoidal curve. (B) The TEAC values of Cpcs and BHT calculated from the results of ABTS assay. Data are shown as mean ± SD. Similar results were obtained from other experiments.

In the CUPRAC assay, which measures SET-based cupric ion (Cu^2+^) reducing capacity. Trolox equivalent (μM TE) of Cpcs exhibited higher activity at lower concentrations than BHT (Fig 3A). The TEAC value of Cpcs exhibited 3.6358 Trolox equivalents per 1 μmol, approximately 2.0-fold higher than that of BHT (Fig 3B).

**Fig 3.**
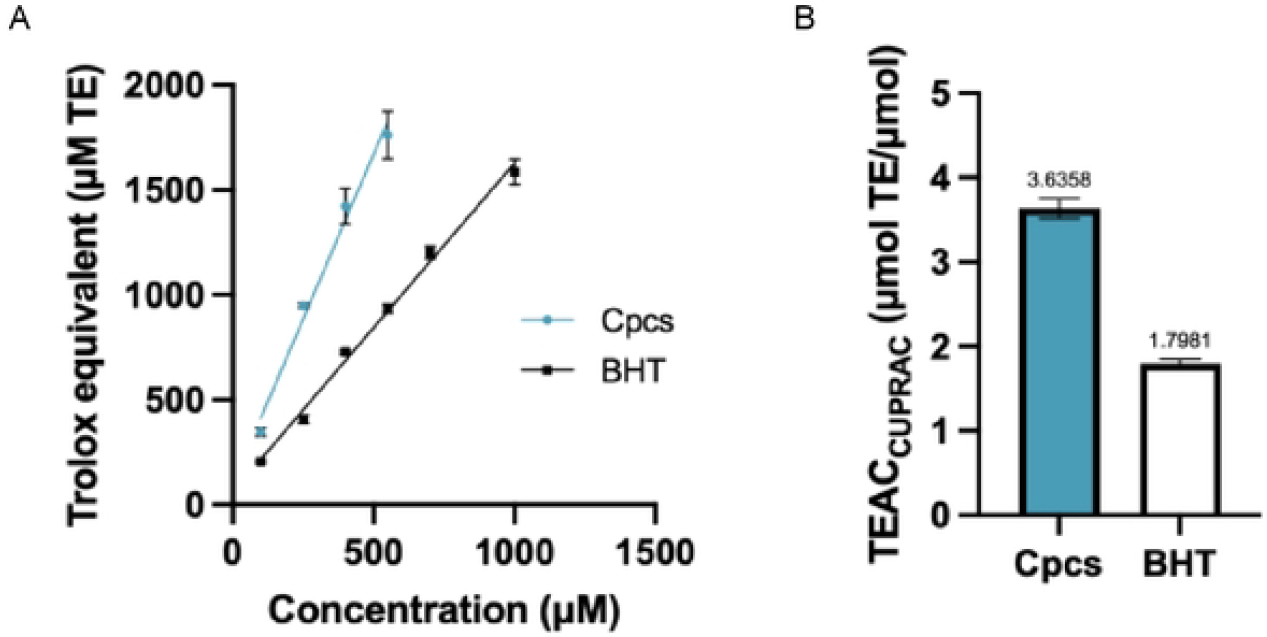
Comparisons of antioxidant capacity between Cpcs and BHT using the CUPRAC assay. (A) Trolox equivalent (μM TE) calculated from the results of CUPRAC assay in 100–1000 μM of Cpcs and BHT. (B) The TEAC values of Cpcs and BHT calculated from the results of CUPRAC assay. Data are shown as mean ± SD. Similar results were obtained from other experiments.

In the ORAC assay, which assesses antioxidant capacity based on peroxyl radical scavenging activity. In the time-dependent decay curves of fluorescence intensity, Cpcs prolonged the decay time in a concentration-dependent manner relative to the control (Fig. 4A). In the TEAC value of Cpcs exhibited 14.186 Trolox equivalents per 1 μmol, approximately 54.7-fold higher than that of BHT (Fig 4B). Although BHT did not exhibit clear concentration-dependent manner, TEAC values of Cpcs and BHT were calculated using data points that falling within the linear range of the Trolox standard curve.

**Fig 4.**
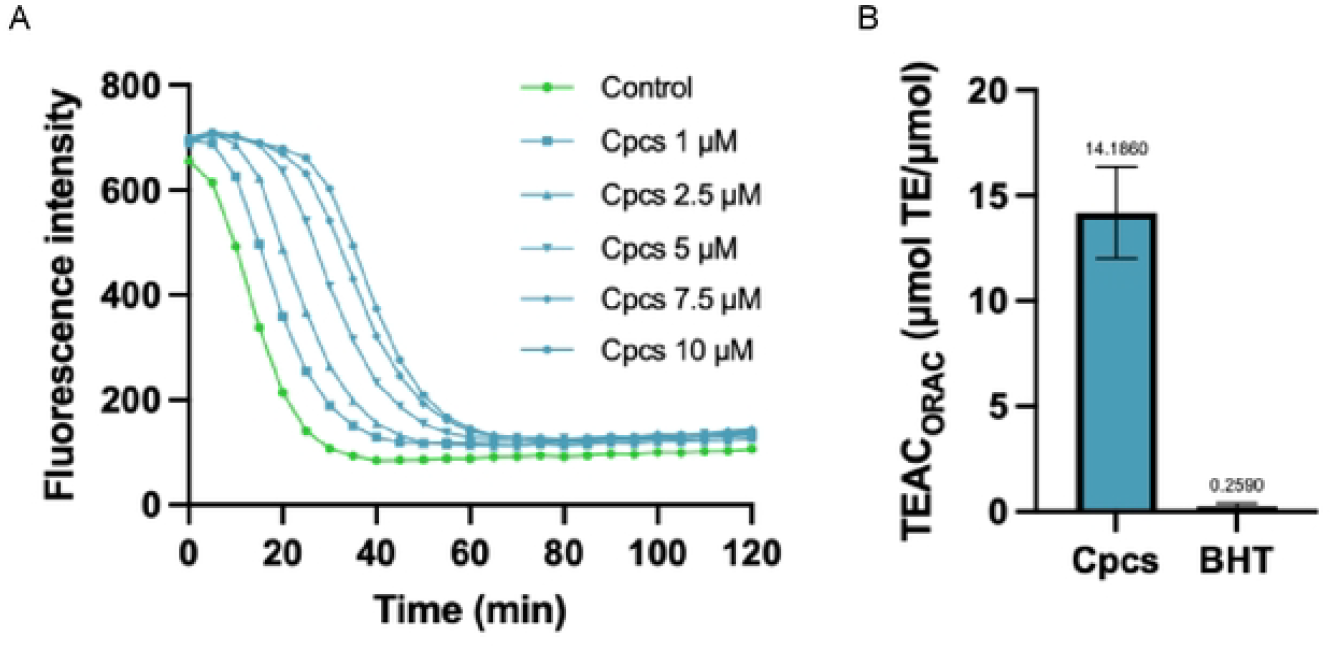
Comparisons of antioxidant capacity between Cpcs and BHT using the ORAC assay. (A) Fluorescence intensities measured by the ORAC assay are shown as the time-dependent decay curves. (B) The TEAC values of Cpcs and BHT calculated from the results of ORAC assay. Data are shown as mean ± SD. Similar results were obtained from other experiments.

### Cpc exhibits cytoprotective activity against rotenone-induced oxidative stress

We examined the cell viability of NHDFs treated with Cpcs at concentrations of 1 or 10 μM. We found that cell viability significantly increased at 10 μM (*p* = 0.0014), indicating that this concentration of Cpcs enhances cell proliferation (Fig 5A). Young NHDFs (passage 5–8) were treated with Cpcs pretreated with Cpcs, and then oxidative stress was induced by using rotenone (Rot), an inhibitor of mitochondrial respiratory complex I. Cytotoxicity was evaluated by measurement of lactate dehydrogenase (LDH) release into culture medium as a marker of cytotoxicity. Cpcs significantly suppressed strong cytotoxicity induced by Rot (*p* = 0.0053) (Fig 5B). Furthermore, since Rot is an inducer of apoptosis [28], cytosolic DNA fragmentation was evaluated by using DAPI staining as an indicator of cell damage related to apoptosis. The results showed that Cpcs significantly suppressed Rot-induced a high degree of DNA fragmentation (*p* = 0.0013) (Fig 5C, D).

**Fig 5.**
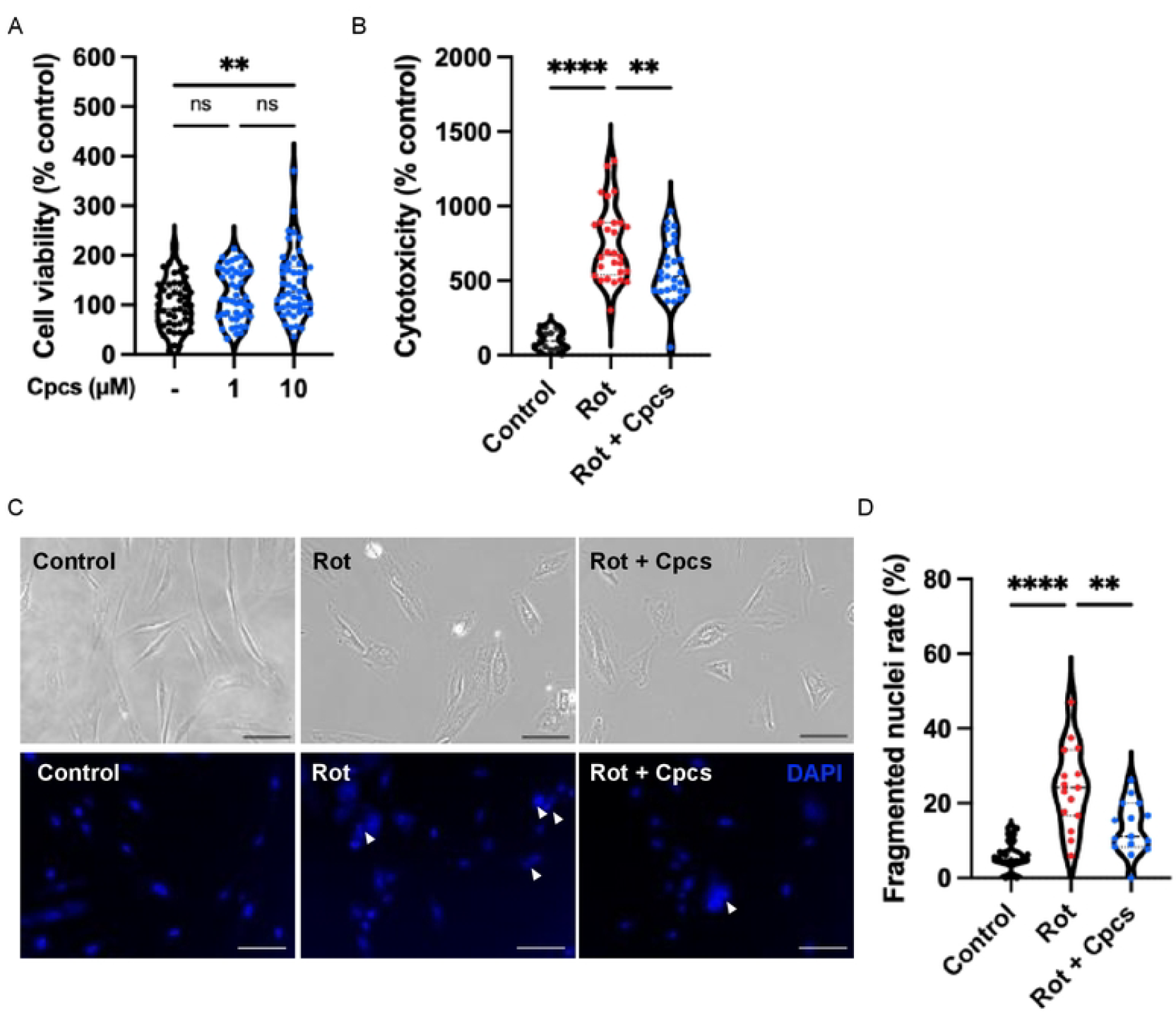
Cytoprotective effects against rotenone-induced oxidative stress by Cpcs. (A) Cell viability in NHDFs treated with 1 or 10 μM of Cpcs (one-way ANOVA with multiple comparisons, \*\**p* < 0.01). (B) Cytotoxicity in NHDFs treated with 1 μM of Rot and 10 μM of Cpcs (one-way ANOVA followed by Tukey’s test, \*\**p* < 0.01, \*\*\*\**p* < 0.0001). (C) Representative microscopy images of cytosolic DNA fragmentation by DAPI staining. White arrow heads indicate fragmented nuclei. Scale bar = 100 μm. (D) Quantification of fragmented nuclei rate treated with 1 μM of Rot and 10 μM of Cpcs (one-way ANOVA followed by Tukey’s test, \**p* < 0.05, \*\*\*\**p* < 0.0001).

## Discussion

In SET-based conditions revealed by the DPPH, ABTS and CUPRAC assays, Cpcs exhibited approximately 2.1–3.6-fold and 2.0–4.3-fold higher antioxidant capacity than Trolox and BHT, respectively. In contrast, Cpcs showed approximately 14.1-fold and 54.7-fold higher antioxidant capacity than Trolox and BHT, respectively, under HAT-based conditions assessed by the ORAC assay. These findings indicate that the antioxidant activity of Cpcs is predominantly governed by the HAT mechanism. This pronounced activity in the HAT-based assay can be explained by the structural features of Cpcs. In HAT reactions, the bond dissociation enthalpy (BDE) of phenolic hydroxyl groups is a key determinant of antioxidant activity [29]. A lower BDE facilitates hydrogen dissociation, and the resulting radical can be stabilized through electron delocalization over the molecular structure [30–31]. The antioxidant capacity of Cpcs exceeded the level expected based solely on the number of hydroxyl groups, suggesting that enhanced radical stabilization significantly contributes to its high capacity.

To elucidate antioxidant property of Cpcs, we compared the TEAC values of Cpcs obtained from this study with those of structurally related compounds that share the catechin or cyanidin skeleton. Cpcs exhibit slightly lower or higher activity trends than flavan-3-ols and comparable or slightly higher activity trends than anthocyanidins and anthocyanins in the SET-based assays (Table 1). In contrast, TEAC values from the ORAC assay indicate that Cpcs surpass representative flavan-3-ols, anthocyanidins and anthocyanins, highlighting its superior performance in the HAT-mediated antioxidant activity as expected (Table 1).

**Table 1.** Comparisons of TEAC with related compounds.

| Compound | TEAC (DPPH) | TEAC (ABTS) | TEAC (ORAC) |
| --- | --- | --- | --- |
| Cpcs | 2.27 ± 0.05 | 2.14 ± 0.18 | 14.1 ± 1.78 |
| Catechin | 0.8 [32] – 2.43 [33] | 1.1 [32] | 7.9 [32] |
| Epicatechin | 1.0 [32] – 2.73 [25] | 1.3 [32] – 1.57 [25] | 5.1 [32] |
| Epigallocatechin | 1.5 [32] – 3.12 [25] | 1.2 [32] – 2.98 [25] | 3.1 [32] |
| Gallocatechin | 8.5 [32] | 4.9 [32] | 8.3 [32] |
| Epigallocatechin gallate | 3.7 [32] – 4.83 [25] | 2.0 [32] – 4.67 [25] | 3.4 [32] |
| Cyanidin | 0.5 [32] | 2.0 [32] | 4.4 [32] |
| Delphinidin | 1.5 [32] | 2.3 [32] | 3.8 [32] |
| Cyanidin-3-*O*-glucoside | 0.6 [32] | 1.9 [32] | 7.3 [32] |
| Cyanidin-3-*O*-galactoside | 0.5 [32] | 2.3 [32] | 5.8 [32] |
| Cyanidin-3-*O*-rutinoside | 0.8 [32] | 1.7 [32] | 5.5 [32] |
| Delphinidin-3-*O*-glucoside | 1.4 [32] | 3.5 [32] | 5.9 [32] |

Data from this study are shown as mean ± SD (Cpcs). The reference values from the literature represent are shown as mean component of values shown as mean (± SD or SEM). All unites are converted to μmol TE/μmol, or the original units (μmol TE/μmol) are cited as is.

The ORAC assay provides a kinetic and biologically relevant evaluation of radical chain antioxidant activity [34–35]. Several of the above-mentioned compounds with established biological activities exhibit high antioxidant capacity based on the HAT mechanism. Cpcs demonstrated greater antioxidant activity than these compounds, suggesting that Cpcs may exert antioxidant capacity in vivo.

In cell culture experiments, Cpcs demonstrated cell proliferation-promoting effects and cytoprotective activity against Rot-induced injury. Rot promotes the production of reactive oxygen species (ROS) by inhibiting mitochondrial respiratory complex I, thereby mitochondria-dependent apoptosis [28, 36–37]. These results suggest that Cpcs may have suppressed the progression of cell death signaling by alleviating oxidative stress.

Since this study suggests that Cpcs may suppress aging and disease through its antioxidant effects, detailed investigations into its in vivo dynamics and physiological activity using cells and animals are required in the future. Moreover, regarding food-industrial applications, since Cpcs are primarily extracted from the seed-coat of beans [21], in traditional Japanese confectionery, for example, Yokan made from boiled adzuki bean paste, the seed coat is sometimes removed during straining and Cpcs are disposed along with the rest. The added value of Cpcs has the potential to promote the effective utilization of such pomace for sustainable health or cosmetic sciences. Therefore, as a new exploration of the useful physiological functions of Cpcs, this research has the potential to yield pioneering results that contribute to a wide range of fields such as nutritional science and food chemistry.

## Acknowledgements

The authors acknowledge the grant from Fukui Prefectural University strategic research promotion grant.

## References

1. Indo HP, Yen H-C, Nakanishi I, Matsumoto K, Tamura M, Nagano Y, et al. A mitochondrial superoxide theory for oxidative stress diseases and aging. J Clin Biochem Nutr. 2014:56(1):1–7. doi: 10.3164/jcbn.14-42.

2. Sakano N, Wang D-H, Takahashi N, Wang B, Sauriasari R, Kanbara S, et al. Oxidative stress biomarkers and lifestyles in Japanese healthy people. J Clin Biochem Nutr. 2009;44(2):185– 195. doi: 10.3164/jcbn.08-252.

3. Tanito M, Takayanagi Y, Ishida A, Ichioka S, Takai Y, Kaidzu S. Liner association between aging and decreased blood thiol antioxidant activity in patients with cataract. J Clin Biochem Nutr. 2023:72(1):54–60. doi: 10.3164/jcbn.22-66.

4. Nanjo F, Mori M, Doto K, Hara Y. Radical scavenging activity of tea catechins and their related compounds. Biosci Biotechnol Biochem. 1999;63(9):1621–1623. doi: 10.1271/bbb.63.1621.

5. Sang S, Cheng X, Stark RE, Rosen RT, Yang CS, Ho H-T. Chemical studies on antioxidant mechanism of tea catechins: analysis of radical reaction products of catechin and epicatechin with 2,2-diphenyl-1-picrylhydrazyl. Bioorg Med Chem. 2002:10(7):2233–2237. doi: 10.1016/s0968-0896(02)00089-5.

6. Anggraini T, Wilma S, Syukri D, Azima F. Total phenolic, anthocyanin, catechins, DPPH redical scavenging activity, and toxicity of Lepisanthes alata (Blume) Leenh. Int J Food Sci. 2019;2019:9703176. doi: 10.1155/2019/9703176.

7. Chen X-Q, Hu T, Han Y, Huang W, Yuan H-B, Zhang Y-T, et al. Preventive effects of catechins on cardiovascular disease. Molecules. 2016;21(12):1759. doi: 10.3390/molecules21121759.

8. Jang H-J, Ridgeway SD, Kim J-A. Effects of the green tea polyphenol epigallocatechin-3-gallate on high-fat diet-induced insulin resistance and endothelial dysfunction. Am J Physiol Endocrinol Metab. 2013;305(12):E1444–1451. doi: 10.1152/ajpendo.00434.2013.

9. Nagao T, Meguro S, Hase T, Otsuka K, Komikado M, Tokimitsu I, et al. A catechin-rich beverage improves obesity and blood glucose control in patients with type 2 diabetes. Obesity. 2008;17(2):310–317. doi: 10.1038/oby.2008.505.

10. Hwang SJ, Yoon WB, Lee O-H, Cha SJ, Kim JD. Radical-scavenging-linked antioxidant activities of extracts from black chokeberry and blueberry cultivated in Korea. Food Chem. 2014;146:71–77. doi: 10.1016/j.foodchem.2013.09.035.

11. Kähkönen MP, Heinonen M. Antioxidant activity of anthocyanins and their aglycons. J Agric Food Chem. 2003;51:628–633. doi: 10.1021/jf025551i.

12. Tsuda T, Watanabe M, Ohshima K, Norinobu S, Choi SW, Kawakishi S, et al. Antioxidative activity of the anthocyanin pigments cyanidin 3-O-.beta.-D-glucoside and cyanidin. J Agric Food Chem. 1994;42(11):2407–2410. doi: 10.1021/jf00047a009.

13. Serra D, Paixão J, Nunes C, Dinis TCP, Almeida LM. Cyanidin-3-glucoside suppresses cytokine-induced inflammatory response in human intestinal cells: Comparison with 5-aminosalicylic acid. PLOS One. 2013;8(9):e73001. doi: 10.1371/journal.pone.0073001.

14. Tang Z. Cyanidin-3-glucoside: Targeting atherosclerosis through gut microbiota and antiinflammation. Front Nutr. 2025;12:1627868. doi: 10.3389/fnut.2025.1627868.

15. Chien C-C, Seiko T, Muto C, Ariga H, Wang Y-C, Chan C-H, et al. A single domestication origin of adzuki bean in Japan and the evolution od domestication genes. Science. 2025;388(6750):abs2871. doi: 10.1126/science.ads2871.

16. Guo Q, Luo J, Zhang X, Zhi J, Yin Z, Zhang J, et al. A comprehensive review of the chemical constituents and functional properties of adzuki beans (*Vigna angulariz*). J Agric Food Chem 2025;73(11):6361–6361. doi: 10.1021/acs.jafc.4c12023.

17. Kitano-Okada T, Ito A, Koide A, Nakamura Y, Han K-H, Shimada K, et al. Anti-obesity role of adzuki bean extract containing polyphenols: in vivo and in vitro effects. J Sci Food Agric. 2012;92(13):2644–2651. doi: 10.1002/jsfa.5680.

18. Kim S, Hong J, Jeon R, Kim H-S. Adzuki bean ameliorates hepatic lipogenesis and proinflammatory mediator expression in mice fed a high-cholesterol and high-fat to induce nonalcoholic fatty liver disease. Nutr Res. 2016;36(1):90–100. doi: 10.1016/j.nutres.2015.11.002.

19. Liu R, Zheng Y, Cai Z, Xu B. Saponins and flavonoids from adzuki bean (*Vigna angularis L.*) ameliorate high-fat diet-induced obesity in ICR mice. Front Pharmacol. 2017;8:687. doi: 10.3389/fphar.2017.00687.

20. Oyama K, Kondo T, Shimizu T, Yoshida K. Determination of absolute configuration of photo-degraded catechinopyranocyanidins A by modified Mosher’s method. Chirality. 2020;32(5):556–563. doi: 10.1002/chir.23202.

21. Yoshida K, Nagai N, Ichikawa Y, Goto M, Kazuma K, Oyama K, et al. Structure of two purple pigments, catechinopyranocyanidins A and B from the seed-coat of small red bean, *Vigna angularis*. Sci Rep. 2019;9(1):16716. doi: 10.1038/s41598-019-53406-9.

22. Yoshida K, Takayama Y, Asano T, Kazuma K. Differences in the content of purple pigments, catechinopyranocyanidins A and B, in various adzuki beans, Vigna angularis. Biosci Biotech Biochem. 2023;87(5):525–531. doi: 10.1093/bbb/zbad010.

23. Apak R, Güçlü K, Özyürek M, Karademir SE. Novel total antioxidant capacity index for dietary polyphenols and vitamins C and E, using their cupric ion reducing capability in the presence of neocuproine: CUPRAC method. J Agric Food Chem. 2004;52(26):7970–7981. doi: 10.1021/jf048741x.

24. Prior RL, Wu X, Schaich K. Standardized methods for the determination of antioxidant capacity and phenolics in foods and dietary supplements. J Agric Food Chem. 2005;53(10):4290–4302. doi: 10.1021/jf0502698.

25. Yamauchi R, Fukamizu S, Kohama Y, Shimamura T, Kashiwagi T, Ukeda H, et al. Comparative DPPH and ABTS radical scavenging activity assays for evaluating natural antioxidants as food additives. Nihon Shokuhin Hozo Kagaku Kaishi (Food Preserv Sci). 2014;40(2):55–63 (in Japanese). doi: 10.5891/jafps.40.55.

26. Kamiya Y, Odama M, Mizuguchi A, Murakami S, Ito T. Puerarin blocks the aging phenotype in human dermal fibroblasts. PLOS One. 2021;16(4):e0249367. doi: 10.1371/journal.pone.0249367.

27. Soccio M, Laus MN, Flagella Z, Pastore D. Assessment of antioxidant capacity and putative healthy effects of natural plant products using soybean lipoxygenase-based methods. An overview. Molecules. 2018;23(12):3244. doi: 10.3390/molecules23123244.

28. Siddiqui MA, Ahmad J, Farshori NN, Saquib Q, Jahan S, Kashyap MP, et al. Rotenone-induced oxidative stress and apoptosis in human liver HepG2 cells. Mol Cell Biochem. 2013;384(1–2):59–69. doi: 10.1007/s11010-013-1781-9.

29. Leopoldini M, Russo N, Toscano M. The molecular basis of working mechanism of natural polyphenolic antioxidants. Food Chem. 2011;125(2):288–306. doi: 10.1016/j.foodchem.2010.08.012.

30. Leopoldini M, Marino T, Russo N, Toscano M. Antioxidant properties of phenolic compounds: H-atom versus electron transfer mechanism. J Phys Chem A. 2004;108(22):4916–4922. doi: 10.1021/jp037247d.

31. Leopoldini M, Pitarch IP, Russo N, Toscano M. Structure, conformation, and electronic properties of apigenin, luteolin, and taxifolin antioxidants. A first principle theoretical study. J Phys Chem A. 2004;108(1):92–96. doi: 10.1021/jp035901j.

32. Tabart J, Kevers C, Pincemail J, Defraigne J-O, Dommes J. Comparative antioxidant capacities of phenolic compounds measured by various tests. Food Chem. 2009;113(4):1226–1233. doi: 10.1016/j,foodchem.2008.08.013.

33. Yamauchi M, Kitamura Y, Nagano H, Kawatsu J, Gotoh H. DPPH measurements and structure-activity relationship studies on the antioxidant capacity of phenols. Antioxidants. 2024;13(3):309. doi: 10.3390/antiox13030309.

34. Knez E, Kadac-Czapska K, Grembecka M. Evaluation of spectrophotometric methods for assessing antioxidant potential in plant food samples – A critical approach. Appl Sci. 2025;15(11):5925. doi: 10.3390/app15115925.

35. Prior RL. Oxygen radical absorbance capacity (ORAC): New horizons in relating dietary antioxidants/bioactives and health benefits. J funct Foods. 2015;18(Part B):797–810. doi: 10.1016/j.jff.2014.12.018.

36. Swarnkar S, Goswami P, Kamat PK, Gupta S, Patro IK, Singh S, et al. Rotenone-induced apoptosis and role of calcium: a study on Neuro-2a cells. Arch Toxicol. 2012;86(9):1387– 1397. doi: 10.1007/s00204-012-0853-z.

37. Li N, Ragheb K, Lawler G, Sturgis J, Rajwa B, Melendez JA, et al. Mitochondrial complex I inhibitor rotenone induced apoptosis through enhancing mitochondrial reactive oxygen species production. J Biol Chem. 2003;278(10):8516–8525. doi: 10.1074/jbc.M210432200.

